# The clinical utility of two high-throughput 16S rRNA gene sequencing workflows for taxonomic assignment of unidentifiable bacterial pathogens in MALDI-TOF MS

**DOI:** 10.1101/2021.08.16.456588

**Authors:** Hiu-Yin Lao, Timothy Ting-Leung Ng, Ryan Yik-Lam Wong, Celia Sze-Ting Wong, Chloe Toi-Mei Chan, Denise Sze-Hang Wong, Lam-Kwong Lee, Stephanie Hoi-Ching Jim, Jake Siu-Lun Leung, Hazel Wing-Hei Lo, Ivan Tak-Fai Wong, Miranda Chong-Yee Yau, Jimmy Yiu-Wing Lam, Alan Ka-Lun Wu, Gilman Kit-Hang Siu

**Affiliations:** Department of Health Technology and Informatics, The Hong Kong Polytechnic University, Hong Kong Special Administrative Region, China; Department of Clinical Pathology, Pamela Youde Nethersole Eastern Hospital, Hong Kong Special Administrative Region, China

## Abstract

Bacterial pathogens that cannot be identified using matrix-assisted laser desorption/ionization time-of-flight mass spectrometry (MALDI-TOF MS) are occasionally encountered in clinical laboratories. The *16S rRNA* gene is often used for sequence-based analysis to identify these bacterial species. Nevertheless, traditional Sanger sequencing is laborious, time-consuming and low-throughput. Here, we compared two commercially available *16S* rRNA gene sequencing tests, which are based on Illumina and Nanopore sequencing technologies, respectively, in their ability to identify the species of 172 clinical isolates that failed to be identified by MALDI-TOF MS. Sequencing data were analyzed by respective built-in analysis programs (MiSeq Reporter Software and Epi2me) and BLAST+ (v2.11.0). Their agreement with Sanger sequencing on species-level identification was determined. Discrepancies were resolved by whole-genome sequencing. The diagnostic accuracy of each workflow was determined using the composite sequencing result as the reference standard. Despite the high base-calling accuracy of Illumina sequencing, we demonstrated that the Nanopore workflow had a comparatively higher taxonomic resolution at the species level. Using built-in analysis algorithms, the concordance of Sanger 16S with the Illumina and Nanopore workflows was 33.14% and 87.79%, respectively. The agreement was 65.70% and 83.14%, respectively, when BLAST+ was used for analysis. Compared with the reference standard, the diagnostic accuracy of optimized Nanopore 16S was 96.36%, which was identical to Sanger 16S and was better than Illumina 16S (71.52%). The turnaround time of the Illumina workflow and the Nanopore workflow was 78h and 8.25h, respectively. The per-sample cost of the Illumina and Nanopore workflows was US$28.5 and US$17.7, respectively.

## INTROUDUCTION

Traditionally, clinical microbiology laboratories have relied on phenotypic methods to identify bacterial pathogens. However, conventional biochemical tests are labor-intensive and time-consuming, and the results can be ambiguous when two species share similar biochemical profiles (1, 2). Nowadays, matrix-assisted laser desorption/ionization time-of-flight mass spectrometry (MALDI-TOF MS) is widely used for bacterial identification in clinical laboratories (3). MALDI-TOF MS allows rapid identification of microorganisms by comparing the mass spectrum of a sample with the reference spectra in the database (4). Although MALDI-TOF MS is a rapid, simple and high-throughput technology for bacterial identification, some species cannot be well differentiated due to high similarity in the mass spectra of closely related species or lack of reference spectra (5).

A study from Lau *et al*. reported that MALDI-TOF MS failed to determine the species of over 70% of phenotypically “difficult-to-identify” bacteria in clinical laboratories(6). In general, anaerobes, particularly *Actinomyces* spp., *Peptostreptococcus* spp., *Prevotella* spp. and *Fusobacterium* spp. (7–9), have a higher failure rate compared with aerobes in bacterial identification using MALDI-TOF MS (7, 10). Additionally, some Gram-positive aerobes, such as *Nocardia* spp. and *Streptomyces* spp., are poorly identified by MALDI-TOF MS (7, 11). Regarding Gram-negative aerobes, studies show that MALDI-TOF MS cannot effectively identify *Acinetobacter* spp., *Chryseobacterium* spp. and *Moraxella* spp. at the species level (11, 12). In such cases, *16S* sequencing of cultured isolates is commonly used for species-level identification.

Sanger sequencing offers a high base-calling accuracy, but it is laborious and time-consuming with limited throughput (13). High-throughput sequencing (HTS) technologies have been proposed as alternatives to generate *16S* sequences for rapid identification of bacteria that are of clinical interest. Next-generation sequencing (NGS), such as can be achieved using Illumina platforms, can generate vast quantities of accurate sequencing reads. However, the read length is limited and insufficient to cover the entire *16S* rRNA gene. According to the official workflow for *16S* rRNA sequencing developed by Illumina Ltd., bacteria are identified based on variable regions (V3 and V4) of *16S*. Nevertheless, these regions are not equally discriminative between and across different species, genera and families (14).

The MinION device by Oxford Nanopore Technologies (ONT) enables generation of reads exceeding 30 kb. The official *16S* rRNA sequencing assay allows the entire *16S* rRNA gene to be sequenced with real-time data analysis. Recent studies have demonstrated its potential for rapid bacterial identification; however, the high read-error rate (8%–15%) of this platform might hinder the accuracy of species-level identification for diagnostic purposes (15).

Considering the respective limitations of Illumina and Nanopore technologies, a comprehensive investigation of the clinical utility of these *16S* rRNA sequencing approaches for bacterial identification is required. This study aimed to evaluate the performance of two commercial HTS workflows for *16S* rRNA sequencing, namely the 16S Metagenomic Sequencing Library Preparation workflow (Nextera XT Index kit v2) from Illumina and the 16S Barcoding Kit 1-24 (SQK-16S024) from ONT, coupled with the respective built-in analysis programs and in-house BLAST+ (v2.11.0) analysis. These workflows were used to identify bacterial isolates that could not be differentiated by MALDI-TOF MS. In light of the complexities of evaluating diagnostic accuracy in the absence of a perfect gold standard, we considered a composite *16S* rRNA sequencing result inferred by Sanger and the two HTS platforms as a reference standard. In case of disagreement in taxa inferred by the three sequencing platforms, whole-genome sequencing (WGS) was conducted to confirm the bacterial identities. In addition, the cost and time-to-result of the sequencing workflows were also compared.

## MATERIALS AND METHODS

### Sample collection and preparation

A total of 172 clinical isolates from 117 species were collected from the clinical microbiology laboratory of Pamela Youde Nethersole Eastern Hospital. Clinical isolates were included if they failed to be classified at the species level (score < 2.00) by the IVD MALDI Biotyper (Bruker Daltonics, Bremen, Germany). Failure to identify bacterial species occurred due to (i) lack of a reference spectrum in the database (81 samples); (ii) inclusion of certain species in the “dangerous database,” named Security Library 1.0, rather than the regular database (two samples); or (iii) poor-quality samples (89 samples) (Table S1). The IVD MALDI Biotyper used in this study was microflex^®^ (Bruker Daltonics), and the database version was BD-6763.

Total nucleic acid was extracted from clinical isolates using the AMPLICOR^®^ Respiratory Specimen Preparation Kit (Roche, Basel, Switzerland) and purified with 1.8X AMPure XP beads (Beckman Coulter, California, USA). Purified DNA was diluted to targeted concentrations in subsequent sequencing workflows. The required DNA input for the Illumina and Nanopore workflows was 12.5 ng and 10 ng, respectively.

### Sanger *16S* rRNA sequencing (Sanger *16S*)

The full-length *16S* rRNA gene was amplified using primers for 16s_008F (5’-AGAGTTTGATCMTGGC-3’) and 16s_1507R (5’-TACCTTGTTACGACTT-3’) (16). The reaction mixture was prepared by mixing 36.7 μl of nuclease-free water, 5 μl of 10× polymerase chain reaction (PCR) buffer, 1 μl of 10-mM deoxynucleoside triphosphate mix (NEB, Ipswich, Massachusetts, USA), 1 μl of each 25-μM primer, 0.3 μl of HotStarTaq Plus DNA Polymerase (Qiagen, Hilden, Germany) and 5 μl of DNA template. The PCR conditions were 96°C for 8 min, 37 cycles at 94°C for 1 min, 37°C for 2 min and 72°C for 2 min 30 s, followed by 72°C for 10 min, and a hold step at 4°C. PCR products were purified using ExoSAP-IT reagent (Thermo Fisher Scientific, Waltham, MA, USA) and then passed to the subsequent cycle sequencing using eight sequencing primers (17–19) (Table S2). The reaction mixture consisted of 13 μl of nuclease-free water, 1 μl of BigDye^®^ Terminator v3.1 Ready Reaction Mix (Thermo Fisher Scientific), 3.5 μl of 5× sequencing buffer, 1 μl of 3.2-μM primer and 1.5 μl of purified PCR product. The PCR conditions were 96°C for 1 min, 25 cycles at 96°C for 10 sec, 37°C for 30 sec and 60°C for 4 min, followed by a hold step at 4°C. The sequencing products were purified using 75% isopropanol and resuspended in 12 μl of Hi-Di^™^ Formamide (Thermo Fisher Scientific). After loading on the Applied Biosystems^®^ 3130 Genetic Analyzer (Thermo Fisher Scientific), the resulting raw trace files were analyzed using the Staden Package (v2.0.0b11). The consensus sequence of each sample was classified by submitting a Basic Local Alignment Search Tool (BLAST) query against the *16S* ribosomal RNA sequence database.

### Illumina sequencing (NGS *16S*)

#### Library preparation

Libraries were constructed according to the 16S Metagenomic Sequencing Library Preparation workflow from Illumina. Briefly, the *16S* V3 and V4 regions of samples were amplified in the first stage of PCR using the primers suggested in the workflow, which were 16S Amplicon PCR Forward Primer (5’-<u>TCGTCGGCAGCGTCAGATGTGTATAAGAGACAG</u>CCTACGGGNGGCWGCAG-3’) and 16S Amplicon PCR Reverse Primer (5’-<u>GTCTCGTGGGCTCGGAGATGTGTATAAGAGACAG</u>GACTACHVGGGTATCTAATCC-3’). The underlined bases in the primer sequences are the overhang adapter sequences for attachment of the indexed adapters in the second stage of PCR. The size of the amplicon was approximately 460 bp. After a post-PCR clean-up, a unique indexed sequencing adapter was added to each sample using the Nextera XT Index kit v2 (Illumina, San Diego, California, USA). Then, a second post-PCR clean-up was performed, followed by a qualification check of the purified libraries.

#### Quantification and sequencing

The size of each library was measured using the 2100 Bioanalyzer system (Agilent, Santa Clara, CA, USA) and the High Sensitivity DNA kit (Agilent). The quantity of the libraries was measured by real-time PCR using the LightCycler^®^ 480 Instrument II (Roche) and QIAseq^™^ Library Quant Assay Kit (Qiagen). Then, the libraries were diluted to 4 nM and pooled into one tube. After denaturation with 0.2-N NaOH, the pooled library was diluted to 9 pM and spiked with 15% of 9-pM PhiX prepared from PhiX Control Kit v3 (Illumina). The pooled library was then loaded on the MiSeq sequencer (Illumina) for sequencing using MiSeq Reagent Kits v3 (Illumina). The sequencing time was 56 h.

#### On-instrument data analysis

Sequencing data were analyzed using MiSeq Reporter software (v2.6.2.3) (MSR) in the MiSeq system. After selecting the metagenomics workflow, sequencing reads were mapped against reference sequences in the Greengenes database (v13.5, May 2013) (http://greengenes.lbl.gov/) for classification. The classification of reads at seven taxonomic levels from kingdom to species was analyzed in this workflow.

#### Data analysis using NGS_BLAST+

The paired-end reads of each sample were merged using the “make.contigs” command in Mothur (v1.44.3) (20). The reads were filtered using the “screen.seqs” command. Sequences smaller than 400 bp, larger than 500 bp, or with any ambiguous bases were removed. The resulting fasta files were analyzed by BLAST+ (v2.11.0) using an in-house Python script (https://github.com/siupenyau/Pocket_16S/tree/7d3fa9d73a6a35afb47e40e7850cef72b4b91a22). In brief, the reads were aligned to the reference sequences in the *16S* ribosomal RNA database (https://ftp.ncbi.nlm.nih.gov/blast/db/) downloaded from the National Center for Biotechnology Information (NCBI). The percentage identity and percentage query coverage were set at 90%.

### Nanopore sequencing (Nanopore *16S*)

#### Library preparation and sequencing

Library preparation was performed using the 16S Barcoding Kit 1-24 (SQK-16S024) from ONT according to the manufacturer’s protocol. Libraries were quantified using the Qubit 2.0 Fluorometer (Thermo Fisher Scientific) with the Qubit^™^ 1X dsDNA HS Assay Kit (Thermo Fisher Scientific). Then, 24 barcoded libraries were pooled into one tube in equal concentrations. After ligation with the rapid adapter, sequencing was performed using the flow cell FLO-MIN106 R9.4.1 with the MinION sequencer on the MinKNOW platform for approximately 4 h.

#### On-instrument real-time data analysis

During sequencing, the passed fastq files, which had a quality score of >7, were uploaded on the cloud-based data analysis platform Epi2me for analysis. Sequencing reads were aligned to reference sequences in the NCBI 16S bacterial database using the FASTQ 16S workflow (v2020. 04. 06). Regarding the workflow parameters, the minimum QSCORE was set at 7, while the minimum percentage coverage and minimum percentage identity were set at 90%.

#### Data analysis using NanoBLAST+

In addition to Epi2me, sequencing data were analyzed using BLAST+ (v2.11.0), similar to the analysis of NGS data. As each sample generated multiple fastq files in a sequencing run, the fastq files of each sample were first merged into a single fastq file and then converted to a fasta file before being aligned to reference sequences in the database.

#### Data analysis using NanoCLUST

Samples with disagreement between EPI2ME and NanoBLAST+ were further analyzed using another pipeline, NanoCLUST (https://github.com/genomicsITER/NanoCLUST) (21). Unlike Epi2me and NanoBLAST+, NanoCLUST does not classify individual reads in a sample. Instead, NanoCLUST forms clusters of similar reads and classifies the consensus sequence of each cluster.

### Whole genome sequencing (WGS)

Samples with complete discordant taxa, as inferred by Sanger *16S*, NGS *16S* and Nanopore *16S* tests, were subjected to WGS to confirm the definite identities using the ONT platform. Library preparation was performed using the transpose-based rapid barcoding kit (SQK-RBK110.96) according to the manufacturer’s protocol. After pooling and adapter ligation, the library was loaded on the flow cell FLO-MIN106 R9.4.1 and sequenced using the GridION device for 48 h in high-accuracy base calling mode. The passed fastq files were uploaded to Epi2me and analyzed using the WIMP workflow (v2021.03.05).

### De novo assembly for WGS datasets

Sequencing reads of each sample were assembled using Shasta (v0.7.0) (https://github.com/chanzuckerberg/shasta). Sequencing reads were re-aligned to the assembled consensus sequences using minimap2 (v2.17-r941) and samtools (v1.10). Consensus sequences were first polished using MarginPolish (v1.3.dev-5492204) (https://github.com/UCSC-nanopore-cgl/MarginPolish) and then further polished using homopolish (v0.2.1) (https://github.com/ythuang0522/homopolish) (22). To avoid bioinformatic bias in de novo assembly, each sample was also subjected to a second analysis pipeline. In brief, the sequencing reads were assembled using miniasm (v0.3-r179) (https://github.com/lh3/miniasm/releases/tag/v0.3). All-vs-all read self-mapping was performed using minimap2. Raw consensus sequences were then generated using miniasm. After re-alignment of the raw reads to consensus sequences using minimap2, the consensus sequences were polished twice using racon (v1.4.3) (https://github.com/isovic/racon).

The longest polished consensus sequences of each sample were classified using BLAST+ (v2.11.0) with the Prokaryotic RefSeq Genomes database downloaded from the NCBI. The top classified species with both query coverage and percentage identity were reported. The average nucleotide identity (ANI) between the query and best-matched reference genomes was calculated using an ANI calculator (https://www.ezbiocloud.net/tools/ani) (23). ANI >94% indicated that the samples belong to the same species as the best-matched genomes.

### Data and statistical analysis

The top classified taxa obtained from NGS and Nanopore datasets were compared with those inferred by Sanger *16S* using built-in programs and BLAST+ for analysis. Species-level concordance between the HTS and Sanger workflows was calculated. For samples that did not match at the species level, concordance at the genus or family level was determined.

To assess diagnostic accuracy, a composite *16S* rRNA sequencing result of the three sequencing platforms was considered as the reference standard. Identical species obtained by at least two sequencing platforms were considered as reference taxa. For samples with complete discordant species inferred by the three sequencing platforms, WGS was conducted to confirm the reference taxa.

## RESULTS

### Statistics of sequencing reads generated from the NGS and Nanopore workflows

Based on the default analysis of MSR, the NGS platform generated an average of 113,381 reads per sample. After merging the paired-end reads and filtering out unwanted reads with undesired read lengths and ambiguous bases, an average of 68,652 filtered reads per sample was retained for NGS_BLAST+ analysis.

The Nanopore MinKNOW platform generated an average of 51,769 reads (QSCORE ≥ 7) per sample, but an average of 51,419 reads (QSCORE ≥ 7) per sample was analyzed in the FASTQ 16S workflow in Epi2me. The slight difference in the number of average reads per sample was due to using different algorithms in the demultiplexing step between Epi2me and Guppy (MinKNOW). An average of 51,769 reads per sample was analyzed using NanoBLAST+.

The total number of reads and the number of classified reads of each sample on both sequencing platforms are shown in Table S3.

### Taxonomic resolution of sequencing reads

The percentage distribution of classified reads via both sequencing platforms is shown in Figure 1. On average, only 45.74% of the total reads of a sample were successfully classified at the species level by MSR with reference to the Greengenes database. After merging paired-end reads and quality filtering, 94.02% of filtered reads were classified at the species level by NGS_BLAST+ with reference to the NCBI *16S* rRNA database.

**Figure 1.**
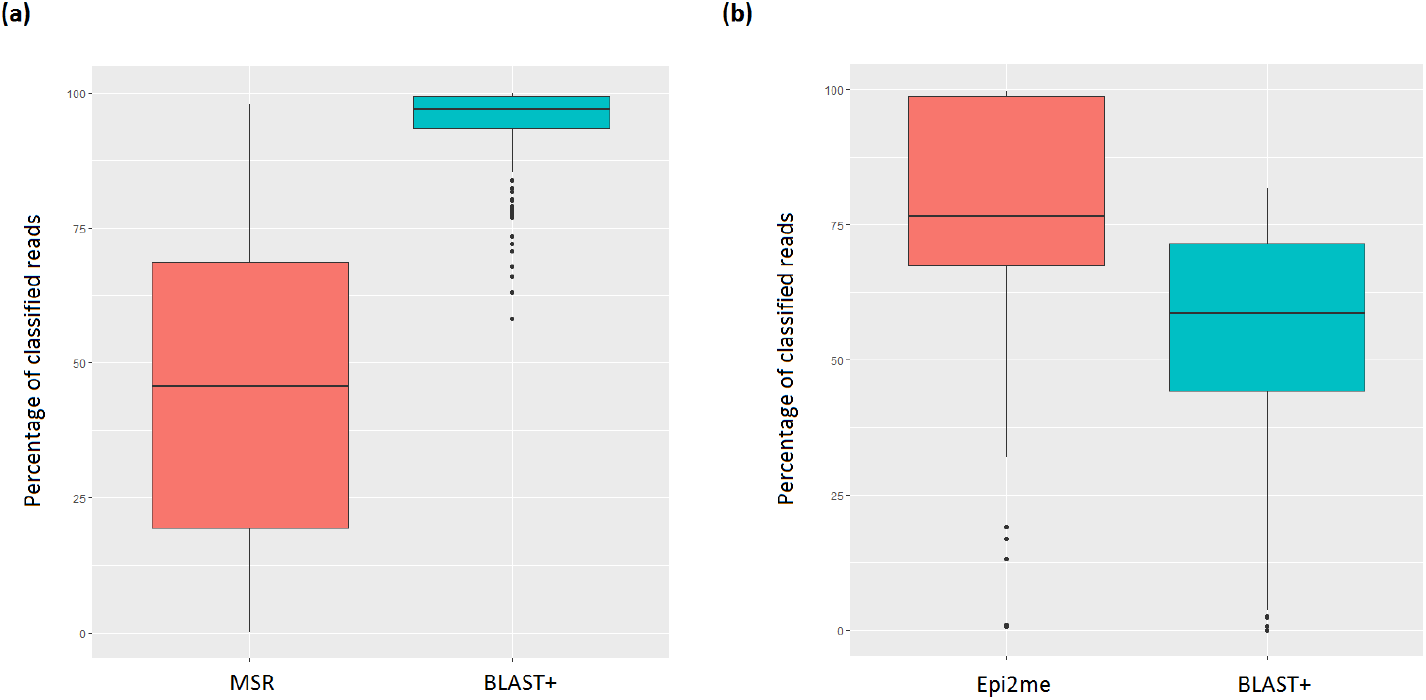
The boxplots showing the distribution of percentage of classified reads of all samples in (a) Illumina and (b) Nanopore sequencing.

In the Nanopore workflow, both Epi2me and NanoBLAST+ use the NCBI *16S* rRNA database for classification of long-read sequencing data. An average of 76.03% of total reads were classified at the species level in Epi2me, compared with 53.56% in NanoBLAST+.

### Concordance in bacterial speciation by Sanger, Illumina and Nanopore *16S* rRNA sequencing

The top-ranked species obtained from the NGS *16S* and Nanopore *16S* workflows, coupled with the respective analysis pipelines, are listed in Table S3 The percentage of samples that matched with Sanger *16S* at each of the species, genus and family levels is illustrated in Figure 2. The concordance in species-level identification among the sequencing platforms is shown in Figure 3. Overall, in terms of concordance with the Sanger *16S* result, Nanopore *16S* was better than NGS *16S* (154/172 [89.53%] vs. 113/172 [65.70%], respectively), regardless of analysis pipeline.

**Figure 2.**
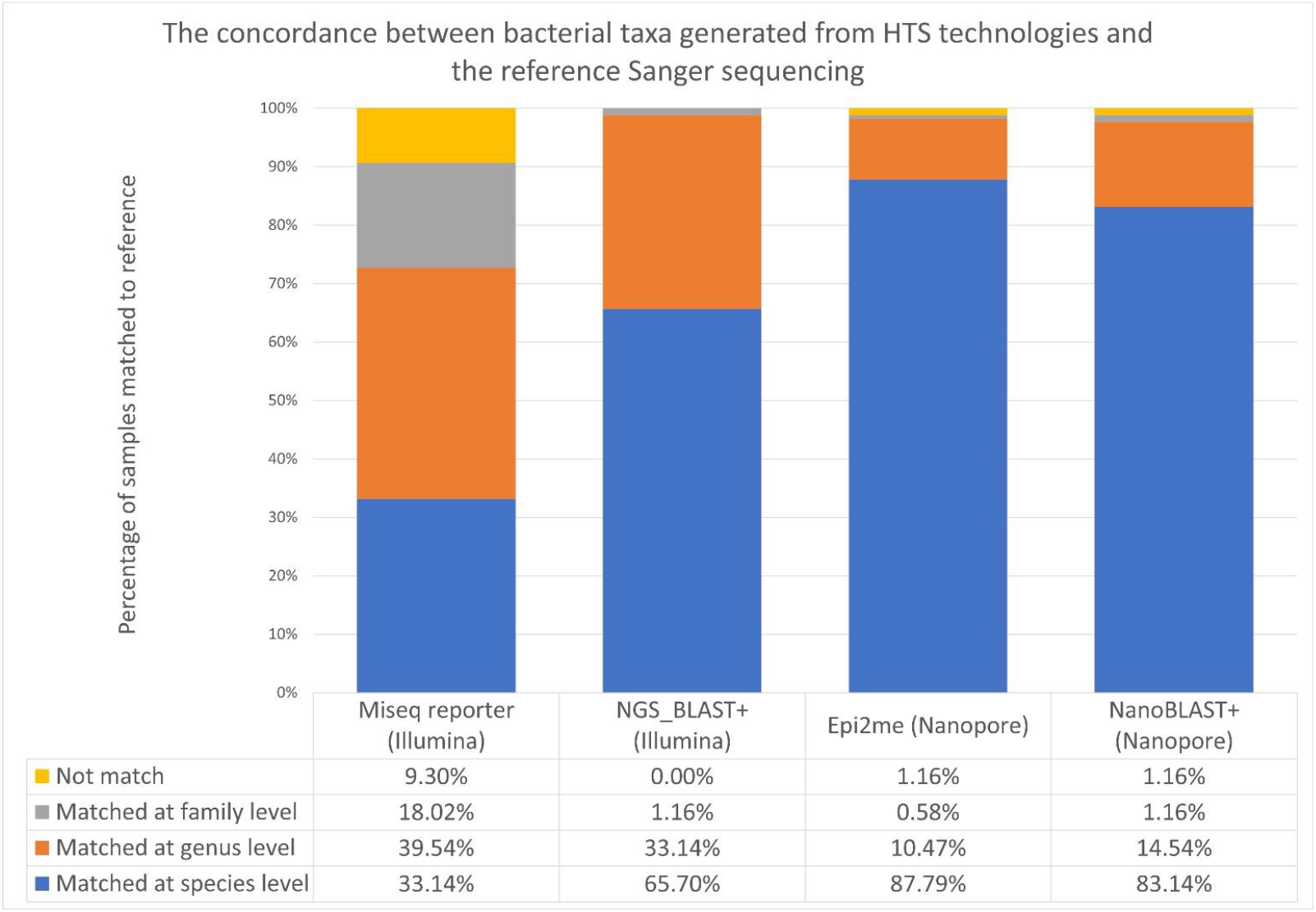
The concordance between bacterial taxa inferred by the two HTS workflows and the Sanger sequencing.

**Figure 3.**
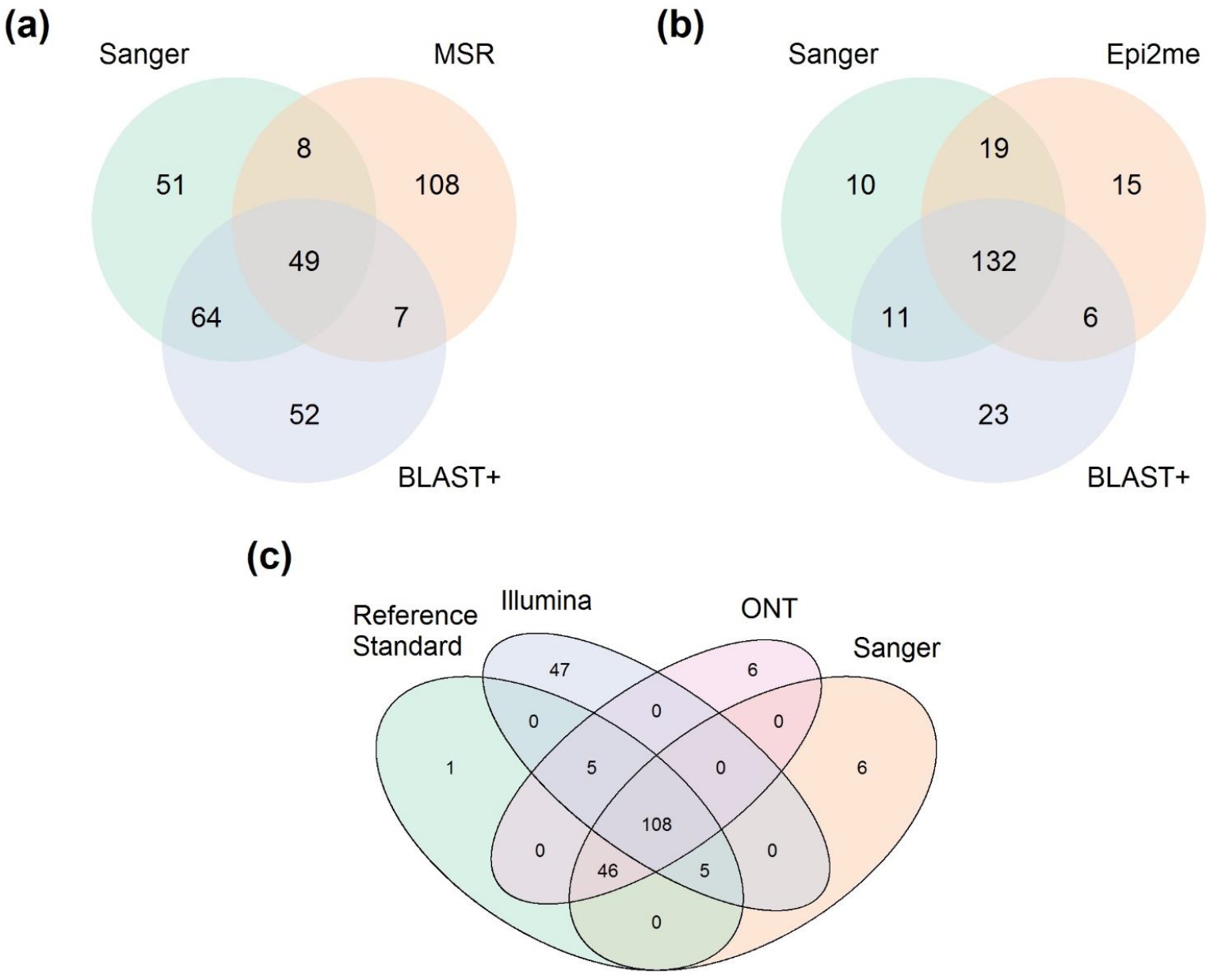
The Venn Diagram showing the concordance of bacterial taxa inferred by different 16S rRNA sequencing platforms. (a) Concordance of top classified species between Illumina sequencing, coupled with MSR and NGS_BLAST+ analysis, and Sanger sequencing. (b) Concordance of top classified species between Nanopore sequencing, coupled with Epi2ME and nanoBLAST+, and Sanger sequencing. (c) Concordance of top classified species among Sanger 16S, NGS 16S, Nanopore 16S and reference standard.

For the NGS *16S* workflow, MSR and NGS_BLAST+ demonstrated a concordance of 33.14% (57/172) and 65.70% (113/172), respectively, with Sanger *16S* in species-level identification. A total of 9.30% of samples (16/172) were unmatched, even at the family level, in MSR, whereas all samples matched at the family level or below in NGS_BLAST+. Of note, concordance between the results of MSR and NGS_BLAST+ was low; only 32.56% of samples (56/172) showed a matched result among the classified species from these two analysis pipelines. Moreover, only 28.49% of samples (49/172) showed complete agreement in the classified species among the MSR, NGS_BLAST+, and Sanger datasets. Owing to poor concordance of the MSR analysis with other sequencing methods, NGS_BLAST+ was considered as the optimal analysis method for the Illumina datasets, and its results were regarded as the final identification inferred by the NGS *16S* workflow.

For Nanopore *16S*, a concordance of 87.79% (151/172) and 83.14% (143/172) at the species level was achieved with Epi2me and NanoBLAST+, respectively. A total of 1.16% of samples (2/172) were unmatched, as reported by Epi2me and NanoBLAST+, respectively. Concordance between the results of Epi2me and NanoBLAST+ was 80.23% (138/172). Additionally, 76.74% of samples (132/172) showed agreement in the classified species among the Epi2me, NanoBLAST+ and Sanger datasets.

A total of 34 samples showed disagreement in the classified species inferred by Epi2me and NanoBLAST+. The respective Nanopore data were further analyzed using NanoCLUST to resolve the discrepancies. NanoCLUST agreed with Epi2ME and BLAST+ in 13 (38.24%) and 17 (50.00%) samples, respectively. Four samples failed to reach agreement in terms of species-level identification, in which three were matched in terms of genus-level identification, and one was considered as having no reliable bacterial ID. Concordance between the resolved Nanopore 16S and Sanger 16S was 89.53% (154/172).

### WGS for bacterial isolates with discrepant species-level ID

Eight samples (4.65% [8/172]) showed complete discordance in bacterial species, as inferred by the three *16S* rRNA sequencing workflows. WGS was conducted to identify definite taxa. Interestingly, seven of these samples failed to match with the published bacterial genomes, with query coverage of <70% for the longest consensus sequences (Table 1). The ANIs to the best-matched genomes were <85% (Threshold for the same species should be >94%), suggesting that these seven “difficult-to-identify” isolates were likely novel bacterial species. As the definite bacterial species could not be confirmed, these samples were excluded from the subsequent diagnostic evaluation.

**Table 1:** Whole genome sequencing analysis for the samples with complete discordant taxonomic assiagnment by Sanger, NGS and Nanopore 16s rRNA sequencing.

| Sample ID | Species inferred by Sanger 16s | Species inferred by NGS 16s | Species inferred by Nanopore 16s | Best-matched Species by WGS (reference genome) | Whole genome sequencing (WGS) |  |  |  |  |  |
| --- | --- | --- | --- | --- | --- | --- | --- | --- | --- | --- |
|  |  |  |  |  | Genome assembly method |  |  |  |  |  |
|  |  |  |  |  | Shasta |  |  | Miniasm |  |  |
|  |  |  |  |  | Query coverage (%) | Identity (%) | ANI (%)<sup>b</sup> | Query coverage (%) | Identity (%) | ANI (%)<sup>b</sup> |
| *Klebsiella pneumoniae* BAA3079<sup>a</sup> | *Klebsiella pneumoniae* | *Klebsiella pneumoniae* | *Klebsiella pneumoniae* | *Klebsiella pneumoniae* (NC_016845.1) | 99 | 97.00 | **98.92** | 92.13 | 99.40 | **99.14** |
| *Staphylococcus aureus* BAA3114<sup>a</sup> | *Staphylococcus aureus* | *Staphylococcus aureus* | *Staphylococcus aureus* | *Staphylococcus aureus* (NC_007795.1) | 94.06 | 99.95 | **99.30** | 88.39 | 99.92 | **99.23** |
| R001 | *Kocuria koreensis* | *Kocuria massiliensis* | *Kocuria spp.* | *Kocuria massiliensis* (NZ_LT835161.1) | 42.21 | 87.44 | **78.29** | 42.42 | 87.41 | **78.55** |
| R006 | *Kocuria koreensis* | *Kocuria massiliensis* | *Kocuria spp.* | *Kocuria massiliensis* (NZ_LT835161.1) | 43.04 | 79.12 | **78.49** | 42.04 | 87.49 | **78.44** |
| R062 | *Klebsiella grimontii* | *Enterobacter cloacae* | *Yokenella regensburgei* | *Klebsiella michiganensis* (NZ_CP060111.1) | 92.17 | 99.17 | **98.71** | 86.30 | 98.99 | **98.69** |
| R120 | *Brachybacterium conglomeratum* | *Brachybacterium faecium* | *Brachybacterium paraconglomeratum* | *Brachybacterium saurashtrense* (NZ_CP031356.1) | 62.15 | 85.18 | **82.30** | 62.30 | 85.12 | **82.39** |
| R121 | *Schaalia odontolytica* | *Schaalia vaccimaxillae* | *Sphingomonas paucimobilis* | *Schaalia odontolytica* (NZ_CP046315.1) | 6.07 | 78.55 | **70.34** | 6.04 | 78.24 | **70.86** |
| R131 | *Schaalia odontolytica* | *Schaalia vaccimaxillae* | *No reliable ID* | *Schaalia odontolytica* (NZ_CP046315.1) | 6.19 | 82.12 | **71.21** | 6.29 | 78.25 | **71.26** |
| R158 | *Microbacterium ginsengite rrae* | *Microbacterium assamensis* | *Microbacterium foliorum* | *Microbacterium foliorum* (NZ_CP041040.1 ) | 65.41 | 84.52 | **82.24** | 65.21 | 84.51 | **82.15** |
| R181 | *Sphingomonas yabuuchi ae* | *Sphingomonas paucimobilis* | *Sphingomonas sanguinis* | *Sphingomonas hominis* (NZ_JABULH01000000 7.1) | 31.48 | 89.67 | **82.09** | 30.68 | 89.59 | **81.95** |

<sup>a</sup> *Klebsiella pneumoniae* BAA3079 and *Staphylococcus aureus* BAA3114 served as QC sample, which were sequenced and analyzed in parallel with the discordant samples for WGS and bioinformatics analysis.
<sup>b</sup> Average Nucleotide Identity (ANI) > 94% indicated that the samples belong to the same species as the best-matched genomes.

The consensus sequence of one sample (R062) showed an overall query coverage of >92%, with 99.17% identity to *Klebsiella michiganensis* (NZ_CP060111.1). As the ANI achieved 98.71%, *K. michiganensis* was therefore considered as the reference taxon for this sample.

### Diagnostic accuracy of the three *16S* rRNA sequencing workflows

Considering the composite of *16S* rRNA sequencing and WGS results as reference standards, the diagnostic accuracy of Sanger *16S*, NGS *16S* and Nanopore *16S* was 96.36% (159/165), 71.52% (118/165) and 96.36% (159/165), respectively, for species-level identification of “difficult-to-identify” bacterial pathogens (Figure 3). The mismatched samples in at least one of the sequencing methods were listed in Table 2. The diagnostic performance of each sequencing workflow was summarized in Table 3.

**Table 2:** The samples with mismatched taxa inferred by at least one sequencing platform.

| Sample ID | Species-level ID (Reference Standard) | Sanger Sequencing (Sanger 16s) |  | Illumina Sequencing (NGS 16s) |  | Nanopore Sequencing (Nanopore 16s) |  |
| --- | --- | --- | --- | --- | --- | --- | --- |
|  |  | Classified species from Sanger 16s <sup>a</sup> | 16s Identity against the reference (%) | Classified species from NGS 16s <sup>a</sup> | 16s Identity against the reference (%) | Classified species from Nanopore 16s <sup>a</sup> | 16s Identity against the reference (%) |
| R003 | *Pseudoglutamicibacter albus* | <u>*Pseudoglutamicibacter cumminsii*</u> | 99.26% | *Pseudoglutamicibacter albus* | matched | *Pseudoglutamicibacter albus* | matched |
| R013 | *Microbacterium hominis* | *Microbacterium hominis* | matched | <u>*Microbacterium aerolatum*</u> | 97.47% | *Microbacterium hominis* | matched |
| R017 | *Microbacterium hominis* | *Microbacterium hominis* | matched | <u>*Microbacterium aerolatum*</u> | 97.47% | *Microbacterium hominis* | matched |
| R021 | *Microbacterium hominis* | *Microbacterium hominis* | matched | <u>*Microbacterium aerolatum*</u> | 97.47% | *Microbacterium hominis* | matched |
| R024 | *Bacillus idriensis* | *Bacillus idriensis* | matched | *Bacillus idriensis* | matched | <u>*Bacillus indicus*</u> | 97.62% |
| R025 | *Varibaculum cambriense* | *Varibaculum cambriense* | matched | <u>*Varibaculum anthropi*</u> | 98.50% | *Varibaculum cambriense* | matched |
| R026 | *Varibaculum cambriense* | *Varibaculum cambriense* | matched | <u>*Varibaculum anthropi*</u> | 98.50% | *Varibaculum cambriense* | matched |
| R036 | *Corynebacterium lowii* | *Corynebacterium lowii* | matched | <u>*Corynebacterium bovis*</u> | 93.29% | *Corynebacterium lowii* | matched |
| R040 | *Weissella cibaria* | *Weissella cibaria* | matched | <u>*Weissella confusa*</u> | 99.26% | *Weissella cibaria* | matched |
| R043 | *Proteus vulgaris* | *Proteus vulgaris* | matched | <u>*Proteus alimentorum*</u> | 99.64% | *Proteus vulgaris* | matched |
| R045 | *Brucella microti* | *Brucella microti* | matched | <u>*Brucella papionis*</u> | 99.86% | *Brucella microti* | matched |
| R047 | *Proteus cibarius* | *Proteus cibarius* | matched | <u>*Proteus terrae*</u> | 99.65% | *Proteus cibarius* | matched |
| R049 | *Dermacoccus barathri* | *Dermacoccus barathri* | matched | <u>*Dermacoccus profundi*</u> | 99.86% | *Dermacoccus barathri* | matched |
| R052 | *Arcanobacterium wilhelmae* | *Arcanobacterium wilhelmae* | matched | <u>*Arcanobacterium pinnipediorum*</u> | 96.60% | *Arcanobacterium wilhelmae* | matched |
| R053 | *Dermacoccus barathri* | *Dermacoccus barathri* | matched | <u>*Dermacoccus profundi*</u> | 99.86% | *Dermacoccus barathri* | matched |
| R056 | *Corynebacterium simulans* | *Corynebacterium simulans* | matched | <u>*Corynebacterium glutamicum*</u> | 93.74% | *Corynebacterium simulans* | matched |
| R058 | *Corynebacterium mastitidis* | *Corynebacterium mastitidis* | matched | <u>*Corynebacterium tuberculostrictum*</u> | 94.67% | *Corynebacterium mastitidis* | matched |
| R062 | *Klebsiella michiganensis* | <u>*Klebsiella grimontii*</u> | 99.20% | <u>*Enterobacter cloacae*</u> | 97.07% | <u>*Yokenella regensburgei*</u> | 98.56% |
| R063 | *Corynebacterium pilbarens* | *Corynebacterium pilbarens* | matched | <u>*Corynebacterium coyleae*</u> | 98.04% | *Corynebacterium pilbarens* | matched |
| R069 | *Eikenella corrodens* | *Eikenella corrodens* | matched | <u>*Eikenella halliae*</u> | 98.69% | *Eikenella corrodens* | matched |
| R071 | *Corynebacterium xerosis* | <u>*Corynebacterium hansenii*</u> | 99.07% | *Corynebacterium xerosis* | matched | *Corynebacterium xerosis* | matched |
| R072 | *Mycolicibacterium fortuitum* | *Mycolicibacterium fortuitum* | matched | <u>*Mycolicibacterium arcuense*</u> | 98.96% | *Mycolicibacterium fortuitum* | matched |
| R073 | *Tessaracoccus oleiagri* | *Tessaracoccus oleiagri* | matched | <u>*Tessaracoccus flavescens*</u> | 95.95% | *Tessaracoccus oleiagri* | matched |
| R078 | *Vagococcus teuberi* | *Vagococcus teuberi* | matched | <u>*Vagococcus martis*</u> | 99.22% | *Vagococcus teuberi* | matched |
| R079 | *Corynebacterium xerosis* | <u>*Corynebacterium hansenii*</u> | 99.07% | *Corynebacterium xerosis* | matched | *Corynebacterium xerosis* | matched |
| R083 | *Tessaracoccus oleiagri* | *Tessaracoccus oleiagri* | matched | <u>*Tessaracoccus flavescens*</u> | 95.95% | *Tessaracoccus oleiagri* | matched |
| R086 | *Raoultella planticola* | *Raoultella planticola* | matched | *Raoultella planticola* | matched | <u>*Klebsiella aerogenes*</u> | 99.06% |
| R094 | *Corynebacterium xerosis* | <u>*Corynebacterium hansenii*</u> | 99.07% | *Corynebacterium xerosis* | matched | *Corynebacterium xerosis* | matched |
| R096 | *Streptomyces thermodiastaticus* | *Streptomyces thermodiastaticus* | matched | <u>*Streptomyces thermoviolaceus*</u> | 98.86% | *Streptomyces thermodiastaticus* | matched |
| R097 | *Pseudoxanthomonas helianthi* | *Pseudoxanthomonas helianthi* | matched | <u>*Pseudoxanthomonas spadix*</u> | 97.04% | *Pseudoxanthomonas helianthi* | matched |
| R098 | *Brachybacterium huguangmaarense* | *Brachybacterium huguangmaarense* | matched | *Brachybacterium huguangmaarense* | matched | <u>*Brachybacterium nesterenkovi*</u> | 97.84% |
| R104 | *Gordonia sputi* | *Gordonia sputi* | matched | <u>*Gordonia otitidis*</u> | 99.07% | *Gordonia sputi* | matched |
| R105 | *Gordonia sputi* | *Gordonia sputi* | matched | <u>*Gordonia otitidis*</u> | 99.07% | *Gordonia sputi* | matched |
| R108 | *Staphylococcus saccharolyticus* | *Staphylococcus saccharolyticus* | matched | <u>*Staphylococcus epidermidis*</u> | 99.19% | *Staphylococcus saccharolyticus* | matched |
| R112 | *Citrobacter sedlakii* | *Citrobacter sedlakii* | matched | <u>*Citrobacter youngae*</u> | 98.32% | *Citrobacter sedlakii* | matched |
| R116 | *Tsukamurella tyrosinosolvens* | *Tsukamurella tyrosinosolvens* | matched | <u>*Tsukamurella ocularis*</u> | 99.86% | *Tsukamurella tyrosinosolvens* | matched |
| R123 | *Pseudoglutamicibacter albus* | <u>*Pseudoglutamicibacter cumminsii*</u> | 99.26% | *Pseudoglutamicibacter albus* | matched | *Pseudoglutamicibacter albus* | matched |
| R133 | *Nocardia brasiliensis* | *Nocardia brasiliensis* | matched | <u>*Nocardia vulneris*</u> | 99.31% | *Nocardia brasiliensis* | matched |
| R140 | *Moraxella lacunata* | *Moraxella lacunata* | matched | <u>*Moraxella equi*</u> | 99.38% | *Moraxella lacunata* | matched |
| R141 | *Ottowia beijingensis* | *Ottowia beijingensis* | matched | <u>*Brachymonas denitrificans*</u> | 93.33% | *Ottowia beijingensis* | matched |
| R149 | *Ornithinibacillus californiensis* | *Ornithinibacillus californiensis* | matched | <u>*Ornithinibacillus scapharcae*</u> | 98.48% | *Ornithinibacillus californiensis* | matched |
| R151 | *Dermacoccus barathri* | *Dermacoccus barathri* | matched | <u>*Dermacoccus profundi*</u> | 99.86% | *Dermacoccus barathri* | matched |
| R153 | *Corynebacterium mastitidis* | *Corynebacterium mastitidis* | matched | <u>*Corynebacterium tuberculostearicum*</u> | 94.67% | *Corynebacterium mastitidis* | matched |
| R175 | *Corynebacterium pollutisoli* | *Corynebacterium pollutisoli* | matched | <u>*Corynebacterium humireducens*</u> | 98.07% | *Corynebacterium pollutisoli* | matched |
| R176 | *Tsukamurella ocularis* | *Tsukamurella ocularis* | matched | *Tsukamurella ocularis* | matched | <u>*Tsukamurella hominis*</u> | 100.00% |
| R178 | *Acinetobacter soli* | *Acinetobacter soli* | matched | *Acinetobacter soli* | matched | <u>*Acinetobacter lactucae*</u> | 97.82% |
| R179 | *Corynebacterium lipophiloflavum* | *Corynebacterium lipophiloflavum* | matched | <u>*Corynebacterium mycetoides*</u> | 97.16% | *Corynebacterium lipophiloflavum* | matched |
| R180 | *Corynebacterium mastitidis* | *Corynebacterium mastitidis* | matched | <u>*Corynebacterium tuberculostearicum*</u> | 94.67% | *Corynebacterium mastitidis* | matched |
| R182 | *Fusobacterium nucleatum* | *Fusobacterium nucleatum* | matched | <u>*Fusobacterium canifelinum*</u> | 98.34% | *Fusobacterium nucleatum* | matched |
| R183 | *Parabacteroides faecis* | *Parabacteroides faecis* | matched | <u>*Parabacteroides chongii*</u> | 97.15% | *Parabacteroides faecis* | matched |
| R190 | *Bacillus xiamenensis* | *Bacillus xiamenensis* | matched | <u>*Bacillus aerius*</u> | 97.16% | *Bacillus xiamenensis* | matched |
| R192 | *Corynebacterium pilbarensis* | *Corynebacterium pilbarensis* | matched | <u>*Corynebacterium ureicelerivorans*</u> | 98.85% | *Corynebacterium pilbarensis* | matched |
| R204 | *Prevotella scopos* | *Prevotella scopos* | matched | <u>*Prevotella jejuni*</u> | 97.41% | *Prevotella scopos* | matched |
| R205 | *Pasteurella multocida* | *Pasteurella multocida* | matched | <u>*Pasteurella stomatis*</u> | 93.74% | *Pasteurella multocida* | matched |
| R206 | *Staphylococcus cohnii* | *Staphylococcus cohnii* | matched | <u>*Staphylococcus auricularis*</u> | 98.16% | *Staphylococcus cohnii* | matched |
| R208 | *Achromobacter denitrificans* | *Achromobacter denitrificans* | matched | <u>*Achromobacter xylosoxidans*</u> | 99.15% | *Achromobacter denitrificans* | matched |
| R210 | *Bacillus licheniformis* | *Bacillus licheniformis* | matched | <u>*Bacillus pascis*</u> | 97.37% | *Bacillus licheniformis* | matched |

**Table 3:**
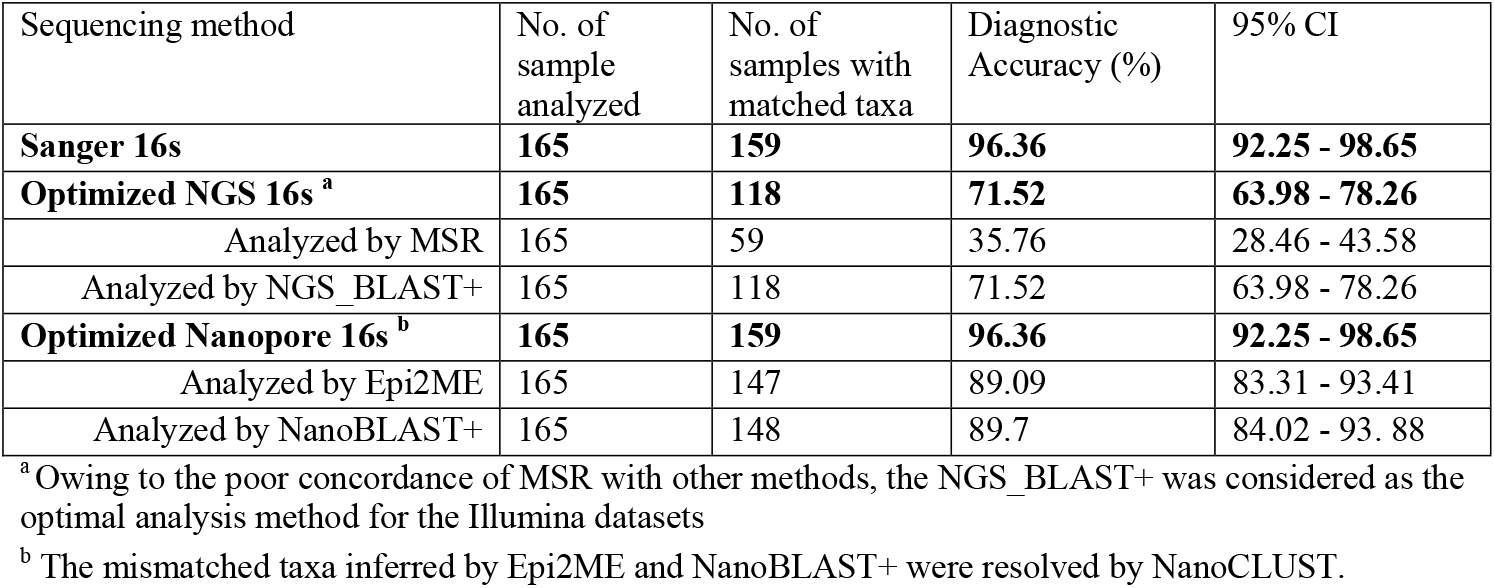
Diagnostic accuracies of the Sanger, NGS and Nanopore 16s rRNA sequencing methods.

### Comparison of sample-to-report time and running cost of the two HTS technologies

The Illumina platform enables sequencing of up to 384 samples per run, whereas, owing to the limited choice of sequencing barcodes, the Nanopore platform can only support a batch of 24 samples per run. Without considering the time for DNA extraction, it took 78 h for the Illumina workflow to generate sequencing data for each run (Figure 4). With the Nanopore platform, the sequencing workflow required 8.25 h. Of note, although base-calling and Epi2me analyses are real-time processes, their speed is highly dependent on the strength of the computer. However, Nanopore sequencing can be stopped once sufficient reads have been generated.

**Figure 4.**
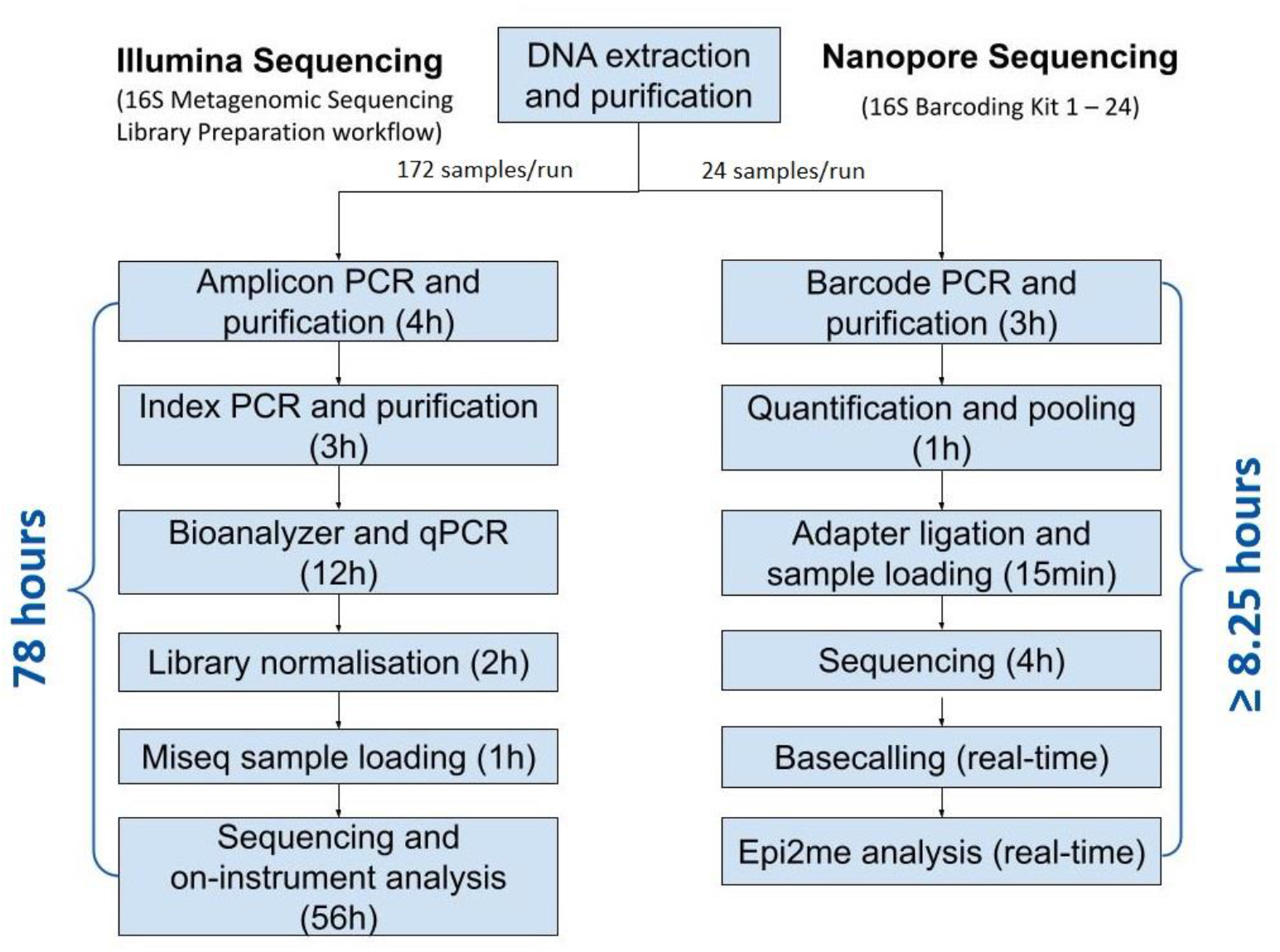
16S rRNA gene sequencing workflow of the HTS technologies.

The running cost of the Nanopore workflow is relatively lower than that of the Illumina workflow. The cost of the Illumina workflow per sequencing run is US $4,931 (172 samples), and the cost per sample is approximately US $28.7. If the sample size is increased to 384, the cost of the Illumina workflow per sequencing run is US $8,279; therefore, the cost per sample is reduced to US $21.6. For the Nanopore workflow, the cost per sequencing run (24 samples) is US $424, which means that the cost per sample is approximately US $17.7.

## DISCUSSION

Although the majority of bacterial pathogens can be identified by MALDI-TOF MS, *16S* rRNA gene sequencing is needed in clinical microbiology laboratories to confirm the identities of “difficult-to-identify” clinical isolates. With reduced costs, simplified protocols and automated bioinformatics pipelines, HTS has been proposed as a better alternative to Sanger sequencing for sequence-based bacterial identification in clinical laboratories. This is the first study to compare the performance (and evaluate the clinical utility) of two commercially available high-throughput *16S* rRNA gene sequencing assays with built-in analysis software for taxonomic assignment of bacterial pathogens that are unidentifiable using MALDI-TOF MS.

With the Illumina platform, the concordance of the classified species between MSR and Sanger *16S* was exceptionally low; only 33.14% of samples matched the reference at top classified species compared with 65.70% when using NGS_BLAST+. As described in previous studies, the use of different bioinformatic tools and *16S* rRNA sequence databases could result in different taxonomic assignments, especially at lower taxonomic levels (24, 25). The latest version of the Greengenes database for MSR was updated in 2013 and does not contain certain new bacterial taxa, which accounts for the poor agreement of this workflow compared with others (25).

Nevertheless, mismatches between NGS and Sanger sequencing were observed in 34.33% of samples, even when the same aligner (i.e., BLAST+) and database (i.e., NCBI 16S bacterial database) were used. One may argue that, with the constraint of low sequencing depth, the Sanger *16S* result alone should not be considered as the final reference. We used a composite of *16S* sequencing results generated by three platforms, and any discrepancies were resolved by WGS as the reference standard to determine the diagnostic accuracy of the HTS workflows. Eventually, a total of 47 samples, including 29 genera and 37 species (Table S3), remained discordant between NGS *16S* and the reference standard. As indicated by Johnson *et al*., although some sub-regions (e.g., V1–V3) of *16S* s rRNA gene provide a reasonable approximation of *16S* diversity, most do not capture sufficient sequence variation to discriminate between closely related taxa. Also, different sub-regions show bias in the bacterial taxa that can be identified (26). In this study, V3–V4 regions might perform poorly in classifying the genera of discordant samples.

Availability of third-generation technologies means that it is becoming possible to exploit the full discriminatory potential of the entire *16S* rRNA gene in a high-throughput manner. The Nanopore *16S* workflow demonstrated a considerably higher percentage concordance with the Sanger *16S* workflow compared with the NGS *16S* workflow, regardless of the analysis pipeline used. In contrast to the built-in analysis on the Illumina platform (i.e., MSR), the performance of Epi2me with Nanopore *16S* was comparable to that of nanoBLAST+ (83.14%), with 87.79% of samples matching Sanger *16S* at top classified species.

Notably, species-level disagreement between Epi2me and nanoBLAST+ was observed in 34 samples (19.77%) and was subsequently resolved by NanoCLUST. Epi2me and BLAST+ rely on read-by-read alignment to reference sequences in the database. As the base-calling accuracy of Nanopore sequencing is relatively low, the prevalence of sequencing errors in Nanopore reads could limit its ability to resolve highly similar sequences. Alternatively, NanoCLUST generates clusters based on Uniform Manifold Approximation and Projection and classifies the representative consensus read in each cluster using BLAST. The effect of sequencing errors in individual sequences can be minimized by forming clusters, which reduces the chance of misclassification. Comparing the species resolved using NanoCLUST with the reference standard, there was a slight improvement in diagnostic accuracy from 89.09% (Epi2me) and 89.70% (nanoBLAST+) to 96.36%.

Six samples (3.64%) failed to match the reference at the species level in the optimized Nanopore *16S* workflow. One possible reason for this discordance is the high similarity in *16S* rRNA gene sequences between the inferred species and the reference taxa. Based on the now historic assumption of *16S* rRNA sequencing, sequences with >95% identity represent the same genus, whereas sequences with >97% identity represent closely related species (27). Many researchers have reported that the taxonomic resolution of *16S* rRNA gene is lower and is unable to discriminate the closely related species in certain genera, including but not limited to *Bacillus*, *Burkholderia*, *Acinetobacter baumannii-calcoaceticus complex*, *Achromobacter*, *Actinomyces* and *Staphylococcus* and the Enterobacteriaceae family (28, 29). In this study, all six taxa inferred by Nanopore *16S* had >97% sequence identity with the reference standard (Table 2).

In this study, WGS was performed to identify the definite bacterial taxa for samples with completely discordant *16S* results. To validate the transposase-based rapid sequencing protocol for bacterial genome construction, two reference strains, namely *Klebsiella pneumoniae* BAA3079 and *Staphylococcus aureus* BAA3114, were sequenced and analyzed in parallel with the eight discordant samples. Both strains successfully yielded consensus sequences of >3Mb, which covered 94% of the genomes of the respective target organisms with 99% identity. This indicated that the WGS protocol was able to construct reliable consensus prokaryotic genomes (Table 1). Nonetheless, the longest consensus sequences of the seven discordant samples failed to obtain a query coverage >50% when mapped to the NCBI Prokaryotic RefSeq Genomes database, suggesting no significant matches between these samples and published bacterial genomes. The ANIs to the best-matched genomes were <94%. These “difficult-to-identify” isolates were therefore considered as novel bacterial species (30). WGS confirmed that R062 belonged to *K. michiganensis* (ANI = 98.71%), which shared a high degree of *16S* rRNA identity with the taxa assigned by Sanger *16S* (*Klebsiella grimontii*; 99.20%), NGS *16S* (*Enterobacter cloacae*; 97.07%) and Nanopore *16S* (*Yokenella regensburgei*; 98.56%) (Table 1). This explains why *16S* rRNA sequencing was not able to accurately differentiate these species.

Considering the time-to-result of the two sequencing platforms, the Nanopore workflow has a much shorter turnaround time compared with the Illumina workflow (8.25 h and 78 h, respectively). Therefore, faster results can be obtained with the Nanopore workflow. However, the sample size is limited to 24 samples per batch. Comparing the cost per sample in a sequencing run, Nanopore sequencing is relatively cheaper than Illumina sequencing (US $17.7 vs. US $28.6, respectively). Additionally, the startup cost of Nanopore sequencing is remarkably lower than that of Illumina sequencing. The starter package of Nanopore sequencing costs only US $1,000, whereas Illumina MiSeq costs approximately US $125,000.

The reusable flow cell FLO-MIN106 R9.4.1, which enables sequencing for up to 72 h, was used for Nanopore 16S in this study. However, library carry over from previous run was observed in a pilot study. This is problematic when the same barcode set is used in consecutive sequencing run. To avoid contamination by library carry over, a new flow cell was used in each sequencing run, and used flow cells were reserved for other sequencing runs using different barcodes. In this context, the disposable Flongle flow cell from ONT is more suitable in a clinical setting. The Flongle flow cell, which costs only US $90, can sequence for up to 16 h. Although the number of active pores available in the Flongle flow cell is lower, it is more cost- and time-effective when the sample size is small. Since it takes time to accumulate a batch of 24 “difficult-to-identify” isolates in clinical laboratories, a small sample size per sequencing run will be beneficial, especially for cases that require urgent diagnosis.

There are some limitations in this study that should be noted. First, the aim of this study was to compare commercially available kits for *16S* rRNA gene sequencing from Illumina and Nanopore. Therefore, by using the 16S Metagenomic Sequencing Library Preparation kit, only the V3–V4 sub-regions of *16S* rRNA gene were sequenced in the Illumina workflow. But it is possible to sequence full-length *16S* rRNA gene using Ilumina MiSeq with a laboratory developed protocol(31), which may increase the taxonomic resolution of the Illumina workflow at the species level. Second, except for the eight discordant samples, the reference taxa of isolates were defined by *16S* rRNA sequencing without being confirmed by WGS. However, some closely related species may have identical *16S* rRNA genes; thus, *16S* rRNA sequencing results may not represent the definite taxa of these samples. Third, regarding the eight samples that underwent WGS, the taxonomic assignment was based on the contigs of consensus sequences after de novo assembly. Circular, gap-free bacterial genomes were not constructed. Finally, bacterial DNA for *16S* sequencing was extracted from cultured isolates. The performance of the NGS *16S* and Nanopore *16S* workflows on direct bacterial identification in microbial and polymicrobial specimens was not evaluated.

## CONCLUSION

In conclusion, the commercial *16S* rRNA gene sequencing workflow from ONT (SQK-16S024), coupled with NanoCLUST, is the most accurate for bacterial identification in a clinical setting, with higher flexibility in sample size and sequencing time, a lower running cost, and higher concordance with the reference standard.

## ACKNOWLEDGMENTS

This work was support by the Innovation and Technology Fund - Partnership Research Programme (PRP) (PRP/010/20FX). The funders had no role in study design, data collection and interpretation, or the decision to submit the work for publication.

